# Genomic variation during culture-adaptation of genetically complex *Plasmodium falciparum* clinical isolates

**DOI:** 10.1101/2022.09.14.507918

**Authors:** Antoine Claessens, Lindsay B. Stewart, Eleanor Drury, Ambroise D. Ahouidi, Alfred Amambua-Ngwa, Mahamadou Diakite, Dominic P. Kwiatkowski, Gordon A. Awandare, David J. Conway

**Author notes:** Corresponding authors: Dr Antoine Claessens; Prof David Conway.

## Abstract

Experimental studies on the biology of malaria parasites have been mostly based on laboratory-adapted lines, but there is limited understanding of how these may differ from parasites in natural infections. Loss-of-function mutants have previously been shown to emerge during culture of some *Plasmodium falciparum* clinical isolates, in analyses that focused on single-genotype infections. The present study included a broader array of isolates, mostly representing multiple-genotype infections which are more typical in areas where malaria is highly endemic. Genome sequence data from multiple time points during several months of culture adaptation of 28 West African isolates were analysed, including previously available sequences along with new genome sequences from additional isolates and timepoints. Some genetically complex isolates eventually became fixed over time to single surviving genotypes in culture, whereas others retained diversity although proportions of genotypes varied over time. Drug-resistance allele frequencies did not show overall directional changes, suggesting that resistance-associated costs are not the main causes of fitness differences among parasites in culture. Loss-of-function mutants emerged during culture in several of the multiple-genotype isolates, affecting genes (including *AP2-HS, EPAC* and *SRPK1*) for which loss-of-function mutants were previously seen to emerge in single-genotype isolates. Parasite clones were derived by limiting dilution from six of the isolates, and sequencing identified *de novo* variants not detected in the bulk isolate sequences. Interestingly, most of these were nonsense mutants and frameshifts disrupting the coding sequence of *EPAC*, the gene with the largest number of independent nonsense mutants previously identified in laboratory-adapted lines. Analysis of Identity-By-Descent to explore relatedness among clones revealed co-occurring non-identical sibling parasites, illustrative of the natural genetic structure within parasite populations.

## INTRODUCTION

Understanding the evolution and adaptation of eukaryotic pathogens requires special efforts, beyond those applied to understanding pathogens with smaller genomes or with higher mutation rates. Malaria parasites are highly adaptive to immunity and to chemotherapeutic or preventive interventions, and with haploid genomes of ∼23 Mb over 14 well-characterised chromosomes they are more amenable than most eukaryotes to systematic study of evolution. The species of greatest medical importance is *Plasmodium falciparum*, one of only two malaria parasite species that have yet been successfully cultured continuously in the laboratory. Understanding parasite adaptation and the processes of evolution may be advanced by deeper analysis of this species, including sampling and cultivation of diverse natural isolates that have not previously been adapted to laboratory conditions.

Previous studies on parasite sequence evolution during culture have focused on laboratory-adapted lines [1, 2], or on clinical isolates that each had single genome sequences prior to culture [3, 4]. Analyses of culture adaptation in 12 different clinical isolates from two West African countries have shown that premature stop codon mutants emerged in approximately half of the isolates during several months of culture [3, 4]. Notably, two gene loci were identified as having independent mutant stop codons emerging in multiple different isolates, an *AP2* transcription factor gene on chromosome 13 (PF3D7_1342900), and the *EPAC* gene on chromosome 14 (locus PF3D7_1417400, encoding Rap guanine nucleotide exchange factor).

Stop codon mutants in other genes including *SRPK1* have each only been seen emerging in single isolates so far, although the small number of isolates examined does not preclude that these genes may also be repeatedly affected. The emergence of independent but convergent mutants indicates that loss-of-function in these genes is likely to be adaptive in culture. This is supported by loss-of-function mutants of the *AP2* gene having a phenotypic effect on growth at variable temperatures which suggests heat shock regulation (thus designated as the *AP2-HS* gene) [5], and by the occurrence of multiple independent stop codons in the *EPAC* gene in other long-term culture adapted lines [3].

For detecting emerging mutants, clinical isolates containing single parasite genotypes were previously focused on as being relatively straightforward to analyse [3, 4], although in endemic populations most *P. falciparum* clinical isolates have multiple genotypes co-circulating in the blood [6]. Initial analysis of these more complex isolates during the process of early culture adaptation has indicated that in most cases there is a gradual loss of genomic diversity [4], but analysis of genomic changes and emergence of new mutants in such isolates has not yet been performed. Here, new sequence data from additional isolates and adaptation timepoints are added to those previously obtained, to enable a substantial survey of parasite genome sequences of 28 *P. falciparum* isolates for up to seven months of culture. Isolates with multiple genomes demonstrated a range of changes in composition during culture adaptation, not explained by previously known fitness costs of drug resistance alleles or *de novo* loss-of-function mutations. Individual parasite clones derived during culture revealed additional *de novo* mutations, and co-occurrence of non-identical sibling parasites which is a feature of natural infections [7, 8].

## METHODS

### Clinical *P. falciparum* isolates from malaria patients

Blood samples were collected from *P. falciparum* malaria cases presenting at government health facilities in Ghana, Guinea, Mali and Senegal, for analysis of parasites at multiple time points after introduction to continuous *in vitro* culture. Twenty-four of the clinical isolates were from patients at Navrongo in the Upper East Region of northern Ghana, as described in previous analyses of parasites at three cultured timepoints [4] with new data presented here on parasite sequences pre-culture and at later timepoints in culture. Two of the isolates were from patients at Faranah in Guinea, one isolate was from a patient at Nioro du Sahel in Mali, and one isolate was from a patient in Pikine, Senegal, for each of which the parasite sequences at multiple timepoints in culture are presented here.

For each isolate, up to 5 ml of venous blood was collected into a heparinised vacutainer (BD Biosciences, CA, USA), and approximately half of the blood sample volume was cryopreserved in glycerolyte at -800C, while the remainder was processed to extract DNA from parasites for whole genome sequencing. In samples from Ghana, leukocytes were removed by density gradient centrifugation and passing through Plasmodipur® filters (EuroProxima, Netherlands), while in samples from Guinea, Senegal, and Mali leukocytes were removed by passing through CF11 powder filtration columns, as previously described [9, 10]. All infections analysed here contained *P. falciparum* alone as determined by Giemsa-stained thick film slide microscopy, except for one Ghanaian sample (isolate 290) that also contained *P. malariae* which does not grow in continuous culture.

### Parasite culture

Cryopreserved patient blood samples were transferred by shipment on dry ice to the London School of Hygiene and Tropical Medicine where culture was performed. Samples were thawed from glycerolyte cryopreservation and *P. falciparum* parasites were cultured continuously at 37oC using standard methods [11] as follows (no isolates were pre-cultured before the thawing of cryopreserved blood in the laboratory on day 0 of the study). The average original volume of cells in glycerolyte in each thawed vial was approximately 1ml, which yielded an erythrocyte pellet of at least 250 µl in all cases. Briefly, 12% NaCl (0.5 times the original volume) was added dropwise to the sample while shaking the tube gently. This was left to stand for 5 mins, then 10 times the original volume of 1.6% NaCl was added dropwise to the sample, shaking the tube gently. After centrifugation for 5 min at 500 g, the supernatant was removed and cells were resuspended in the same volume of RPMI 1640 medium containing 0.5% Albumax™ II (Thermo Fisher Scientific, Paisley, United Kingdom). Cells were centrifuged again, supernatant removed and the pellet resuspended at 3% haematocrit in RPMI 1640 medium supplemented with 0.5% Albumax II, under an atmosphere of 5% O2, 5% CO2, and 90% N2, at 370C, with orbital shaking of flasks at 50 revolutions per minute. Replacement of the patients’ erythrocytes in the cultures was achieved by dilution with fresh erythrocytes from anonymous donors every few days, so that after a few weeks of culture parasites were growing virtually exclusively in erythrocytes from new donors. All clinical isolates were cultured in parallel in separate flasks at the same time, so that the donor erythrocyte sources were the same for all the different isolates, enabling comparisons without confounding from heterogeneous erythrocytes.

### Generating parasite clones by limiting dilution

Parasitized erythrocytes were diluted to 2 per ml mixed with uninfected erythrocytes at 1% haemotocrit, and 250µl transferred to individual wells of a 96 well plate (a probability of 0.5 parasites per well). On each plate, 12 wells with uninfected erythrocytes at 1% haematocrit (negative control) and 12 wells initially having 100 parasitized erythrocytes per well at 1% haematocrit (positive control) were used to monitor the presence of parasites by microscopy. The plate was gassed in a culture chamber with 5% CO2, 5% Oxygen and 90% Nitrogen and incubated at 370C. The medium in each well was replaced after 24h with fresh complete medium and subsequently at Day 4, 7, 10 and 14. Fresh erythrocytes were added at Day 4 and all wells were diluted 5-fold at Day 14. 50µl of positive and negative control wells were removed on Day 11 for PCR targeting the serine tRNA ligase gene locus (PF3D7_0717700). The erythrocytes were pelleted by centrifugation and the blood pellet added to the PCR reaction mixture in a final volume of 5µl using KAPA Blood PCR kit (peqlab) which does not require a separate DNA extraction, using the recommended cycling conditions in the kit. At Day 11, negative control wells remained PCR negative whilst all positive control wells were positive by PCR. At Day 21, 50µl was removed from each of the cloning wells and PCR was performed as described above.

PCR positive wells at this point were assumed to contain parasite clones and culture of each of these was scaled up into 5ml volumes at 1% haematocrit.

### Genome sequencing of parasites at different timepoints in culture

DNA extracted from parasites at each of the assayed culture timepoints and each clone was used for whole-genome Illumina short-read sequencing. Library preparation, sequencing and quality control was performed following internal protocols at the Wellcome Sanger Institute, similarly to previous sequence generation from clinical isolates from these populations [6, 9, 10]. As part of the process, a protocol to enrich parasite genomic DNA compared to human DNA by selective whole genome amplification does not have a significant effect on the within-sample parasite sequence composition and diversity [12]. Genetic variants were called using a pipeline developed by the MalariaGEN consortium (ftp://ngs.sanger.ac.uk/production/malaria/Resource/28), with short reads mapped to the *P. falciparum* 3D7 reference genome sequence [13] version 3 using the BWA algorithm. Single Nucleotide Polymorphisms (SNPs) and short insertions-deletions (indels) were called using GATK’s Best Practices. Variants with a VQSLOD score < 0 or within the large subtelomeric gene families *var, rif* and *stevor* were excluded, to focus on reliable scoring variants in the core genome.

### Within-isolate genomic diversity, emergence of new mutants and analysis of relatedness among clones

Estimation of parasite diversity within each isolate relative to overall local population diversity was performed using the *F*WS index [14, 15] (a within-isolate fixation index between 0 and 1.0 that is inverse to the level of diversity such that a value of 1.0 indicates a pure clone), as previously applied to analysis of some of the cultured timepoints of the Ghanaian clinical isolates [4]. Analysis of new mutants followed methods similar to those previously used in analysis of clinical isolates with single genotype infections [3, 4], except that isolates with mixed genotype infections were analysed in this study. Allele frequencies of intragenic SNPs and frameshift-causing indels covered by read depths of at least 10 were plotted for each timepoint, to scan for the emergence of new mutant alleles. The quality of the mapped sequence reads for each of the cases of putative mutants were inspected visually using the Savant software [16]. Although SNP calling proved straightforward in mixed infection isolates, indel calling was not generally reliable due to the mixed signal, so indels in single genotype isolates and clones were focused on. Finally, to estimate genetic relatedness between genomes of different parasite clones, Identity-By-Descent (IBD) was calculated using hmmIBD using default parameters [17].

## RESULTS

### Whole genome sequencing of *P. falciparum ex vivo* and after up to 7 months of culture adaptation

Analysis of parasite genome sequence variation was performed on 28 *P. falciparum* clinical isolates from West Africa, grown in culture for periods of several months (range 76 to 204 days, median 153 days). An aliquot of the culture of each isolate was sampled for Illumina whole-genome sequencing on multiple occasions (a median of three, and up to five timepoints) (Fig. 1). Overall, 96 high-quality genome sequences from these uncloned parasite cultures were investigated here (Supplementary Table S1), of which 67 were previously generated in a study of Ghanaian isolates that analysed sequences of single-genotype isolates, but which did not investigate novel variants within the multiple-genotype isolates [4]. The 29 new uncloned parasite genome sequences here comprised 17 additional timepoints for the Ghanaian isolates (including 12 at day zero prior to culture) and three timepoints from each of four isolates from other countries (Fig. 1 and Supplementary Table S1).

**Fig. 1.**
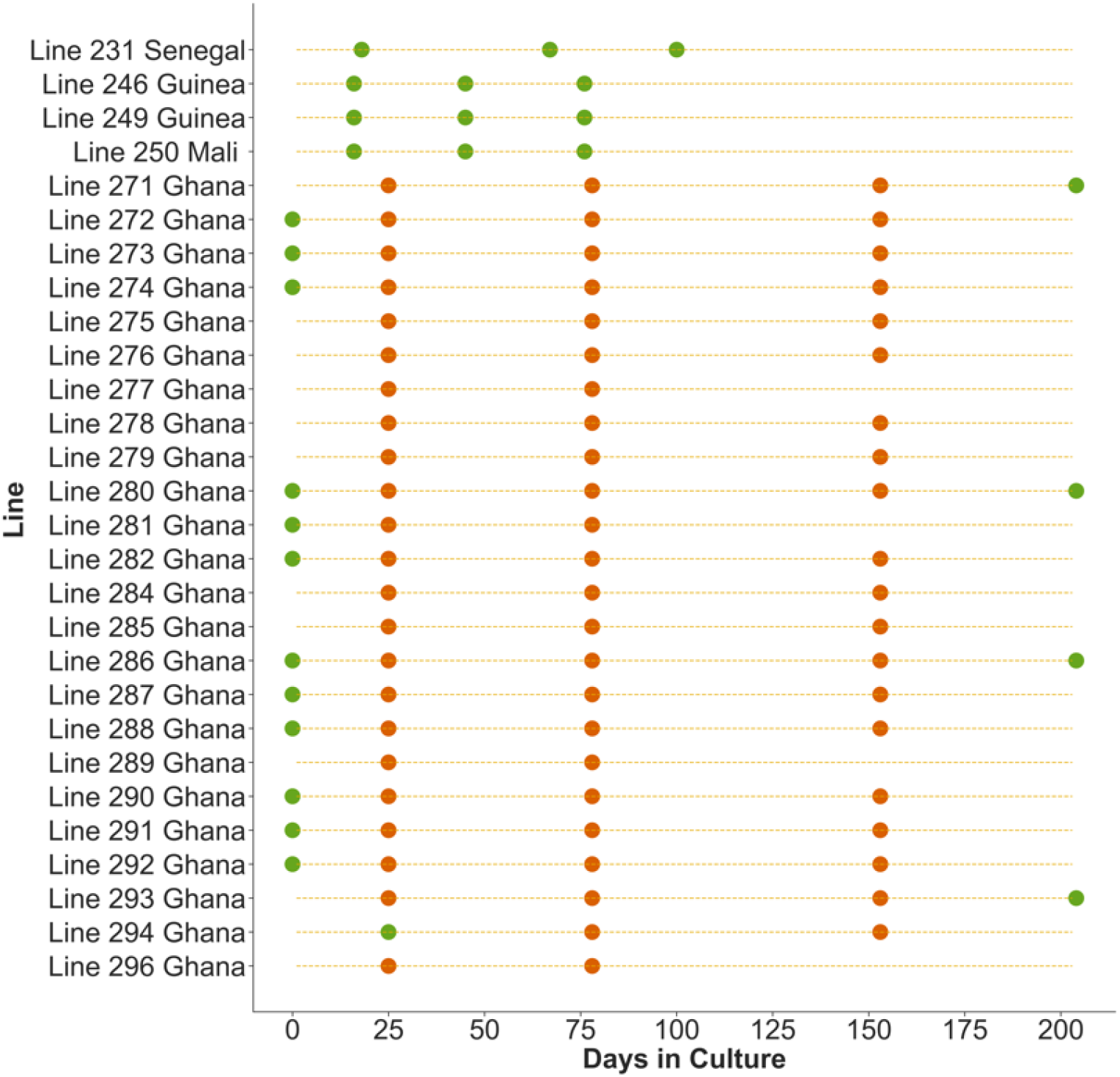
Scheme indicating sampling of genome sequences of *P. falciparum* clinical isolates at different timepoints during the process of culture adaptation. Circles indicate culture time points of 28 isolates sampled for whole genome sequencing. Orange symbols indicate data previously obtained in a study of Ghanaian isolates, and green symbols indicate timepoints for which sequences were derived in the current study. The new data include additional timepoints for 15 of the Ghanaian isolates (including a pre-culture sample for 12 of these) as well as data for four isolates from other West African countries. Sequence accession numbers are given in Supplementary Table S1.

A summary measure of the genomic complexity of parasites within isolates was first obtained by calculating the *F*WS fixation index which ranges from 0 to 1 (an isolate with a value > 0.95 having predominantly a single genome while lower values indicate more complex mixtures of different parasite genotypes). The overall trend was that within-isolate genomic diversity declined during culture adaption, as shown by the increasing *F*WS index values over time (Fig 2A & B), as previously noted for the isolates from Ghana [4]. However, in approximately one third of the isolates (9 out of 28), there were periods when the *F*WS index decreased between successive timepoints, with declines of more than 0.1 in the values indicating temporary increases in diversity (Fig 2C). As such patterns could reflect faster growing genotypes being sometimes initially rare within infections, or might reflect more complex processes, the profiles of allele frequency changes were next examined directly for each isolate.

**Fig. 2.**
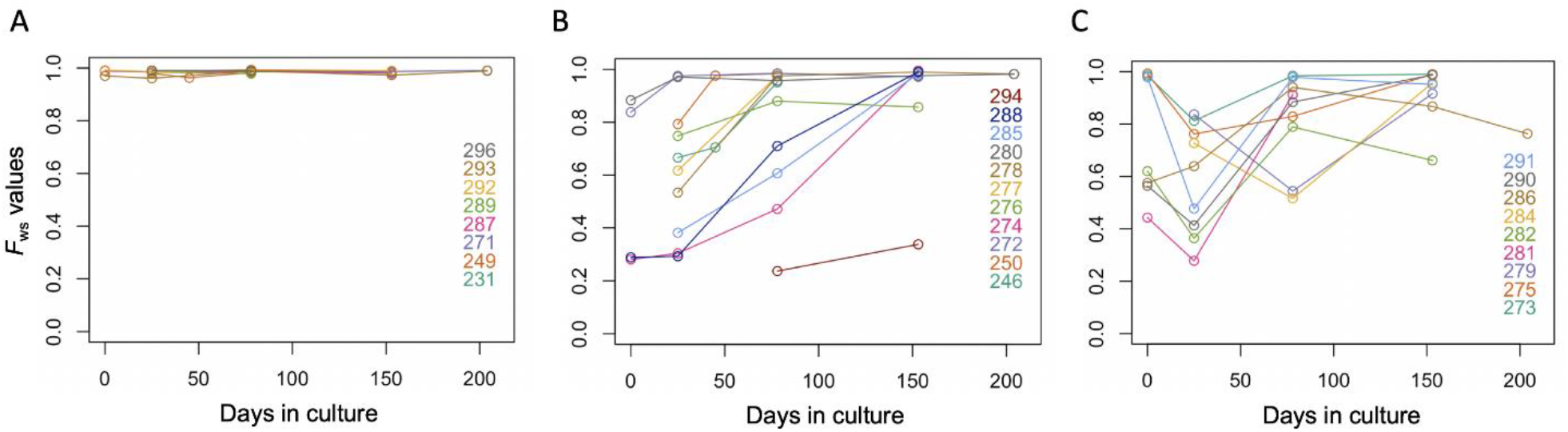
Different patterns of genomic diversity changes during culture of *P. falciparum* clinical isolates. Data for 28 different West African isolates are shown, individual isolates being labelled by identification number and colour. For each timepoint, the within-isolate fixation index *F*WS inversely indicates the genetic complexity, with values close to 1.0 indicating isolates having single genotypes and lower values indicating greater complexity. (A) Eight isolates show single genotypes throughout the culture adaptation period. (B) Eleven isolates each show progressive reduction of genetic diversity over time. (C) Nine isolates show more complex patterns including a temporary increase in genetic diversity (*F*WS index decreasing by a value of at least 0.1) occurring at some point during the culture adaptation period. The *F*WS values for each individual sample timepoint for each isolate are given in Supplementary Table S1.

### Clonal diversity within an isolate varies in different ways during culture adaptation

Allele frequencies of all SNPs were plotted for each isolate at every sample timepoint (Fig. 3 and Supplementary Fig. S1). Consistent with the summary shown by analysis of the *F*WS indices, these plots show that in most isolate lines the genetic diversity gradually reduced over time in culture. 23 parasite lines showed evidence of containing multiple genotypes in at least one of the early timepoints sampled. Of these, 13 had only a single genome detected by the end of the culture period, indicating that some *P. falciparum* genotypes outcompete others. This is apparent from the clearly parallel trajectory of allele frequency changes in most SNPs within the isolates, although the phase of haplotypes is not directly known by bulk sequencing. In some of the isolate lines, the frequency changes over time appear to follow a simple directional pattern (Fig. 3A). However, some of the lines displayed more complex changes in proportions of different genotypes over time (Fig. 3B). For example, the allele frequencies within Line 284 indicate a minority genotype at day 25 that increased to majority at day 77, but then decreased to near disappearance by day 153.

**Fig 3.**
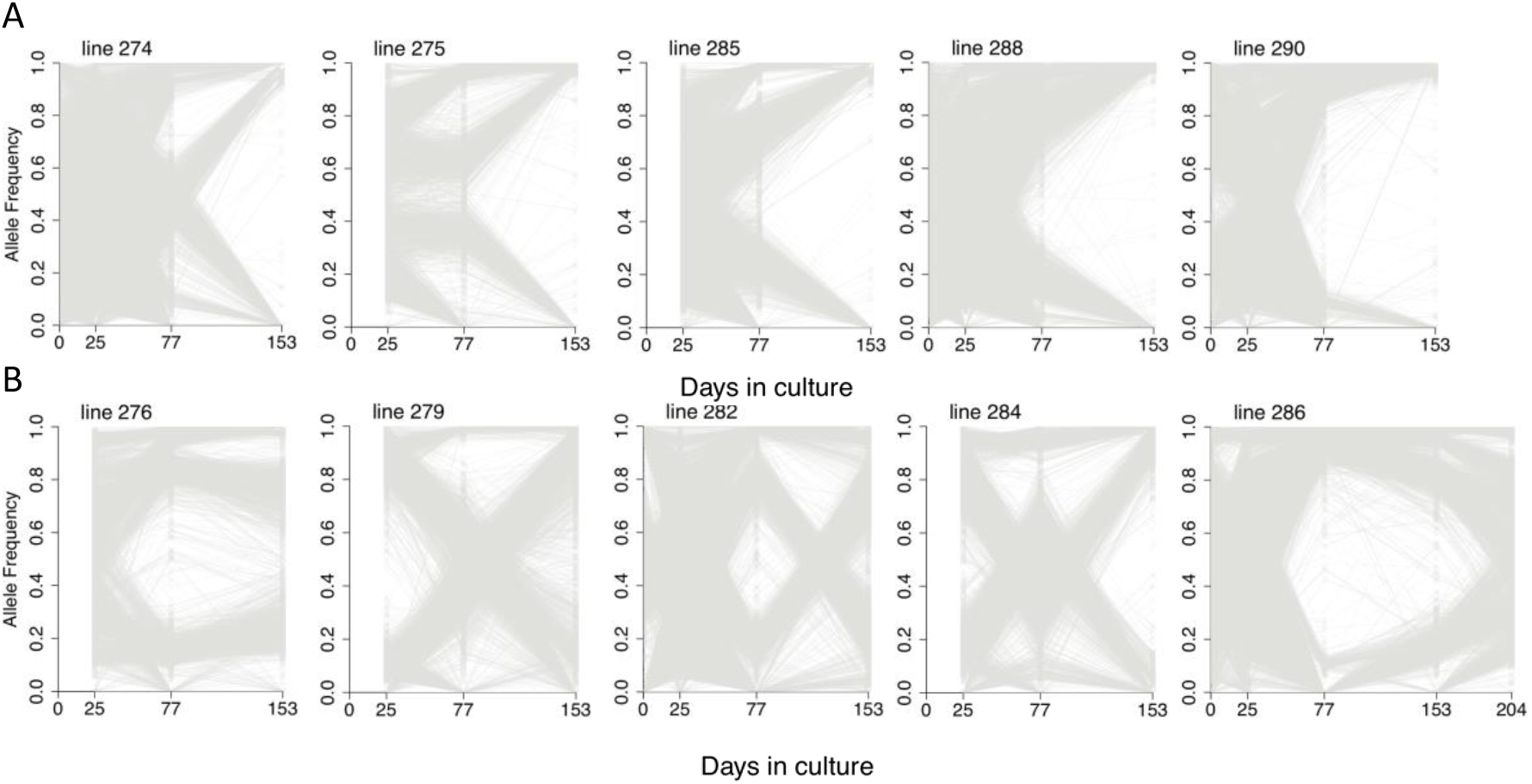
Genome-wide plots of single nucleotide polymorphism allele frequencies within *P. falciparum* multiple-genotype clinical isolates during culture adaptation. Allele frequency of each SNP within an isolate was estimated by the proportion of sequencing reads matching the reference (3D7 genome to which all sequences were mapped) or alternative allele for each nucleotide position. Grey lines indicate allele frequencies changing over time. Data for 10 of the isolates are shown. (A) Five isolates polyclonal at the first sequenced timepoint and monoclonal by day 153. (B) Five isolates polyclonal at the first sequenced timepoint, with variation in the proportion of each genome, that do not reach fixation by the end of the culture adaptation period. Plots for all isolates, including those not illustrated here, are shown in Supplementary Fig. S1.

### Drug resistance alleles do not explain most changes in genotype frequencies in culture

As drug resistance alleles may carry a fitness cost compared to wild-type alleles, we assessed their frequencies within polyclonal isolates during culture adaptation (Fig. 4 and Supplementary Fig. S2). A consistent decrease in resistant allele frequencies over time could indicate that genomes bearing wild-type alleles have a growth advantage. Four genes that have established drug resistance alleles circulating in Africa were examined (*crt, mdr1, dhfr* and *dhps*) with the main drug-resistance-related allele for each gene being shown in Fig. 4 (other variants are shown in Supplementary Fig. S2). The *dhps* codon S436A is a marker of resistance to Sulphadoxine, *dhfr* codon S108N resistance to Pyrimethamine, *crt* K76T resistance to Chloroquine and *mdr1* N86Y resistance to aminoquinolines including Chloroquine. Out of 13 isolates mixed for the *dhps* S436A polymorphism, seven showed a decrease and six an increase in the resistance-associated allele frequency during culture adaptation. For the *dhfr* S108N polymorphism, out of six mixed isolates two showed a decrease and four an increase in the resistance-associated allele frequency. Similarly, no *mdr1* or *crt* drug-resistant allele showed a trend in allele frequency change across the different isolates. Over all four genes, there was no evidence of a significant decrease in drug resistance marker allele frequencies over time, and no overall directionality to allele frequency changes (Binomial test, P > 0.5).

**Fig 4.**
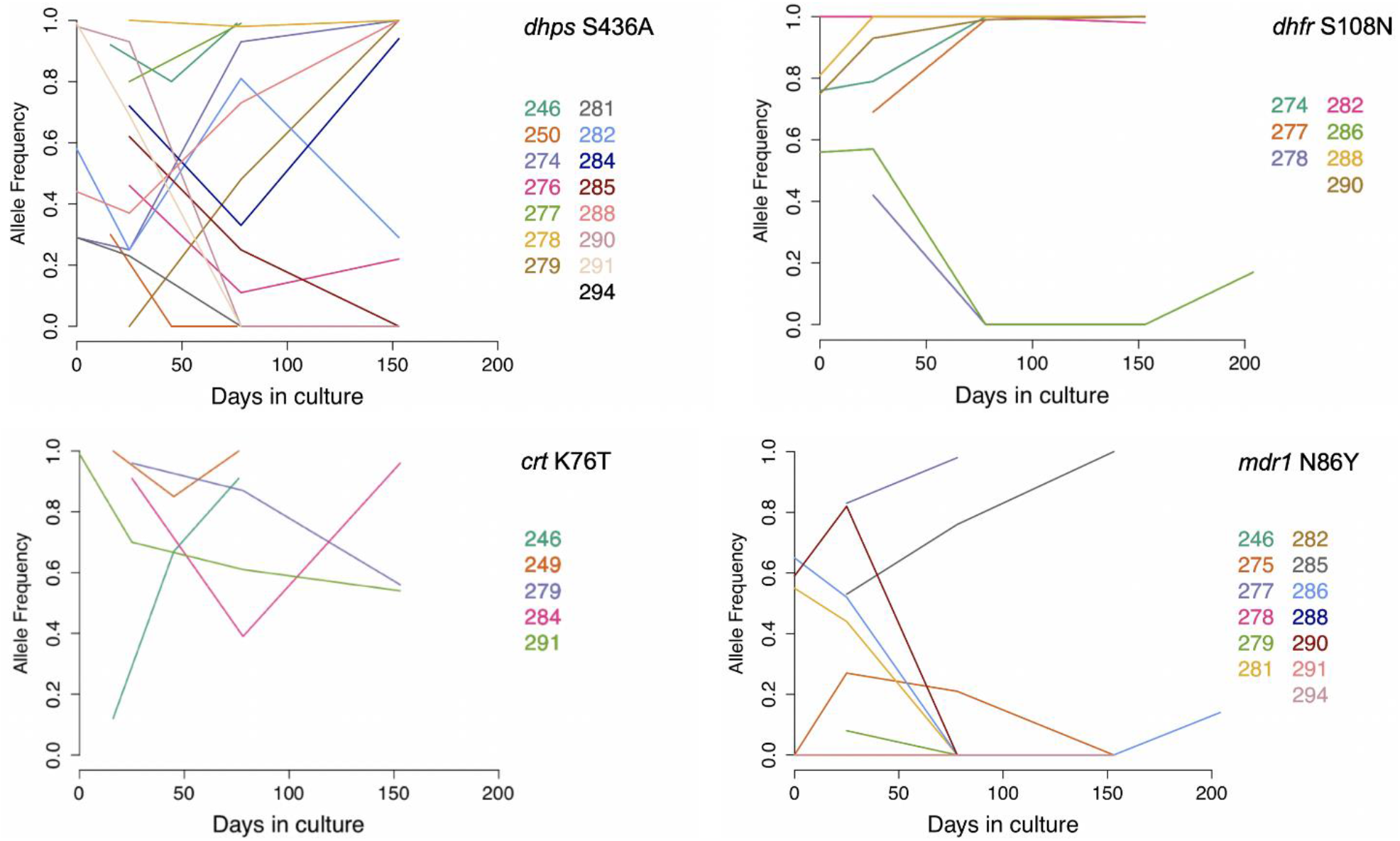
Frequencies of known drug resistance-associated alleles within *P. falciparum* multiple-genotype clinical isolates during several months of culture. Plots show data for one common resistance-associated polymorphism within each of four different genes (chloroquine resistance genes *crt* and *mdr1* on chromosomes 7 and 5, and antifolate resistance genes *dhfr* and *dhps* on chromosomes 4 and 8, respectively). For each of these polymorphisms, coloured lines indicate the data for isolates that had mixed alleles at one or more timepoints. The allele frequencies of other polymorphisms within these genes are shown in Supplementary Fig. S2. Across all isolates, there was no significant directionality to the changes in frequencies of any of these resistance-associated polymorphisms over time in culture.

### Loss-of-function mutations emerging during culture

In previous studies on single genotype infection isolates, loss-of-function mutants appeared to have a selective advantage during culture growth. Here, to examine mixed genotype isolates, analysis was conducted on stop codon mutations and indels generating frameshifts, as either type of mutation would lead to loss of function. Eight isolates showed at least one loss-of-function mutant rising in frequency during culture. Three of these were single-genotype isolates for which we had previously described the mutants [4] and five of which were multiple-genotype isolates for which the mutants were not previously described (Fig. 5 and Table 1).

**Fig 5.**
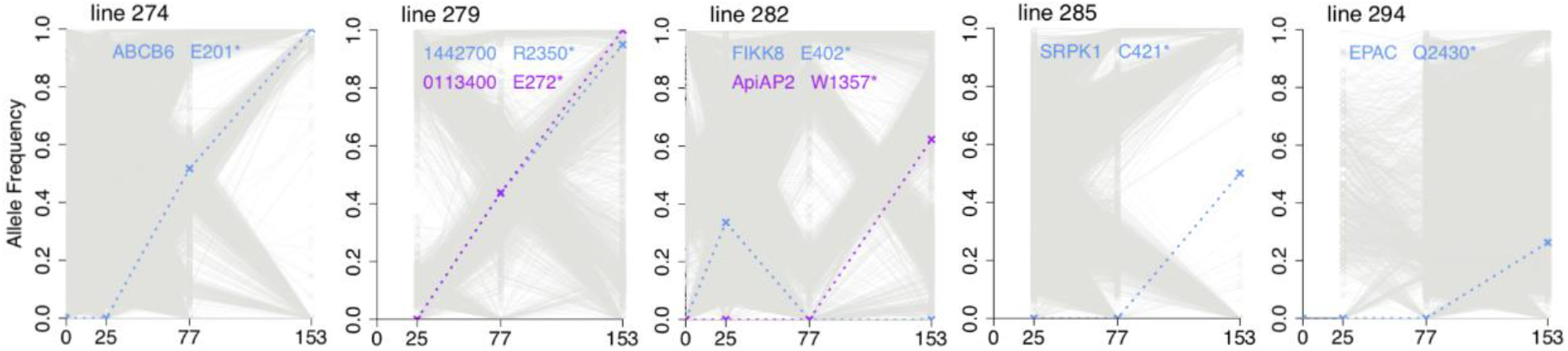
Emergence of premature stop codon mutants during culture of mixed-genotype *P. falciparum* clinical isolates. Stop codon alleles that emerged and attained a frequency of at least 0.2 are shown, with dotted lines and labelling of the gene and codon affected. Grey lines indicate all other SNP allele frequencies changing over time within these isolates. Separate to these new findings from mixed-genotype isolates, emerging mutants in three of the single-genotype isolates (lines 271, 272 and 280) were previously reported [4]. Apart from these, no other isolate among the 28 studied here had a loss-of-function mutant sequence detected to reach a frequency of 0.2 during culture.

**Table 1.**
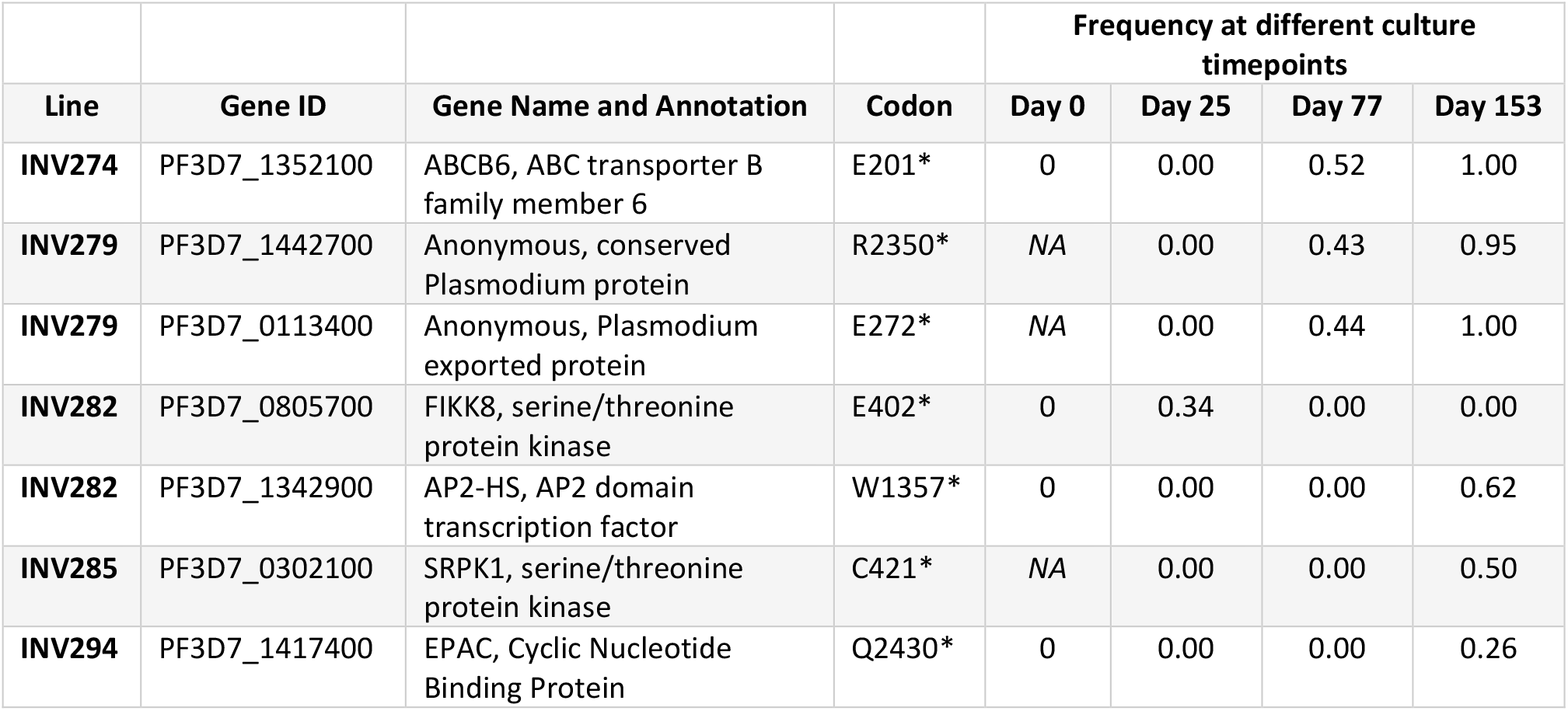
Novel mutants encoding premature stop codons emerging in multiple-clone *P. falciparum* clinical isolates.

| Line | Gene ID | Gene Name and Annotation | Codon | Frequency at different culture timepoints |  |  |  |
| --- | --- | --- | --- | --- | --- | --- | --- |
|  |  |  |  | Day 0 | Day 25 | Day 77 | Day 153 |
| INV274 | PF3D7_1352100 | ABCB6, ABC transporter B family member 6 | E201* | 0 | 0.00 | 0.52 | 1.00 |
| INV279 | PF3D7_1442700 | Anonymous, conserved Plasmodium protein | R2350* | NA | 0.00 | 0.43 | 0.95 |
| INV279 | PF3D7_0113400 | Anonymous, Plasmodium exported protein | E272* | NA | 0.00 | 0.44 | 1.00 |
| INV282 | PF3D7_0805700 | FIKK8, serine/threonine protein kinase | E402* | 0 | 0.34 | 0.00 | 0.00 |
| INV282 | PF3D7_1342900 | AP2-HS, AP2 domain transcription factor | W1357* | 0 | 0.00 | 0.00 | 0.62 |
| INV285 | PF3D7_0302100 | SRPK1, serine/threonine protein kinase | C421* | NA | 0.00 | 0.00 | 0.50 |
| INV294 | PF3D7_1417400 | EPAC, Cyclic Nucleotide Binding Protein | Q2430* | 0 | 0.00 | 0.00 | 0.26 |

Each of these mutants may be considered separately. The genes affected in two of the isolates have not been previously seen to have loss-of-function mutants emerge in culture. In the isolate line 274, a premature stop codon in an ABC transporter gene *ACCB6* (at codon position 201) was associated with the genome sequence of an initial minority parasite that was undetectable at day 25, which replaced parasites with other genomes to become fixed within the culture by day 153. No stop codon is seen in this gene in sequences from several thousand clinical *P. falciparum* infections [6], and the orthologous gene is essential for *in vivo* replication of the rodent malaria parasite *P. berghei* in mice [18]. Interestingly, an insertional mutagenesis screen of the laboratory-adapted *P. falciparum* strain NF54 indicated that disruption of this gene strongly reduced the parasite growth rate in culture [19], which suggests that fitness affects are conditional on the parasite genetic background. In the isolate line 279, there was a replacement of parasites with different genomes over time between days 25 and 153, with stop codon mutations in two different genes being associated with the genotype going to fixation. One of these genes (Pf3D7_0113400) encodes an exported protein [20], while the other encodes a protein of unknown function (Table 1). Insertional mutagenesis has previously indicated both genes as dispensible for the growth of parasites in culture [19]. In one isolate a premature stop codon in a serine/threonine kinase gene *FIKK8* was seen in a minority of sequence reads at day 25 of culture, but not detected in any of the earlier or later timepoints, an observation not replicated or associated with directional change (Table 1).

In isolate lines 282, 285 and 294, premature stop codons were detected respectively in *ApiAP2-HS, SRPK1* and *EPAC* (Fig. 5 and Table 1). In each case, the mutants were not seen until after day 77 of culture and had intermediate frequencies that were still far from fixation by day 153.

Other independent premature stop codon mutations were previously seen to emerge in these same three genes during culture of single-genotype Gambian clinical isolates[3], and two of these genes (*ApiAP2-HS* and *EPAC*) had premature stop codon mutations emerging during culture of the single-genotype Ghanaian isolates [4]. It is likely that a slight growth fitness advantage is conferred by loss-of-function mutation of these genes, as mutants have emerged on different single-genome isolate backgrounds, where they were not associated with other genomic changes, as well as in the mixed-genotype isolates here.

As no loss-of-function mutants were detected during culture of the other isolates analysed here, the overall proportion of isolates with such emerging mutants was 29% (8 out of 28). There was no significant difference in the proportions for single-genotype isolates (38%, 3 out of 8) and multiple-genotype isolates (25%, 5 out of 20)(Fisher’s Exact test, P = 0.65).

### Identification of *de novo* mutations in parasite clones

Bulk whole genome sequencing may only be likely to detect new variants in a parasite under positive selection, as most new mutants will be very rare. To investigate whether other variants may be present, we performed parasite cloning by limiting dilution of six of the clinical isolates after 100 days of culture. The parasite clones were then grown for up to 104 days, and for an initial scan up to 3 of these clones from each isolate were sequenced, generating 17 clone sequences in total (Fig 6A). Novel SNPs or indels were detected in five of the 17 clones, one from each of five different isolates (Table 2). Interestingly, four of these variants are loss-of-function mutations in the *EPAC* gene, a different novel variant in a clone from each of four different isolates. This is further evidence that such mutants commonly arise within the *EPAC* gene, the locus with the most loss-of-function mutants previously detected. The only other locus with novel variant among the parasite clone sequences is an anonymous gene (Pf3D7_320700) in which a nonsynonymous coding change was detected, but it is notable that this gene has been described to have a premature stop codon variant in the long-term culture-adapted line Dd2 [3].

**Fig 6.**
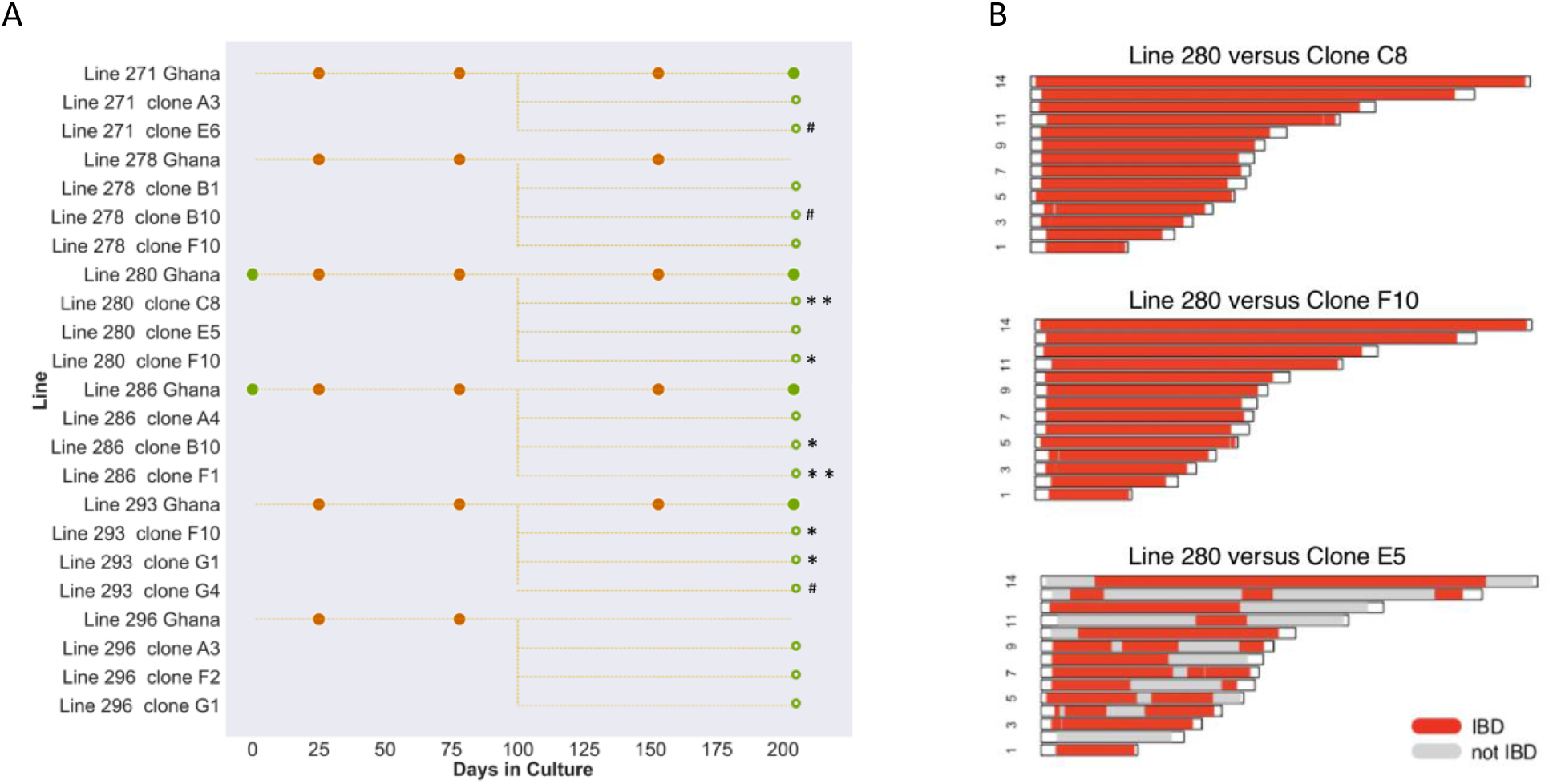
Genome sequencing of parasite clones derived from isolates show additional novel mutants as well as sibling parasites. (A) After 100 days of culture, clones were derived from six of the clinical isolate lines by limiting dilution, and 104 days after the cloning step a few of these clones from each isolate were sampled for sequencing (17 in total), indicated by hollow circles. These sequences were compared with the bulk sequence of the corresponding isolate (at day 77). Clones in which novel SNPs and indels were detected are indicated by asterisks and hashtags respectively (the mutants are detailed in Table 2 and Supplementary Table S1). (B) Clones from each isolate were scanned for genomic patterns of Identity-By-Descent (IBD). The three panels show the IBD in the 14 chromosomes, comparing the bulk cultured Line 280 (at day 77) with three derived clones. Sub-telomeric regions at the end of each chromosome were not analysed and are unshaded. Clone E5 shows 60.8% of the genome in IBD with the Day 77 genome as well as with Clone 1 and 2, consistent with a full sibling relationship. The clones from the other isolates showed complete identity to their respective isolate bulk culture majority sequences, although deeper sampling would likely reveal more cases of non-identical sibling parasites.

**Table 2.**
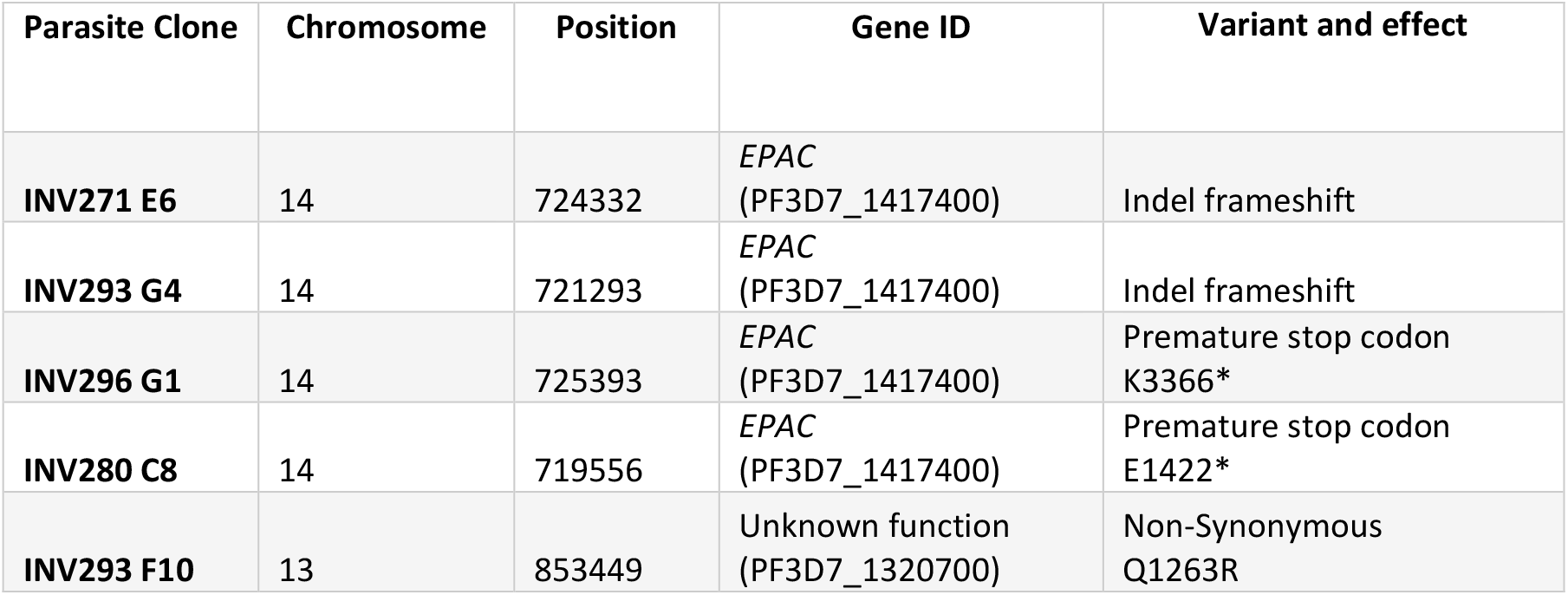
Sequence variants identified in parasites cloned from clinical isolates. Variants also present in the parental isolate, and likely conferring a growth advantage.

| Parasite Clone | Chromosome | Position | Gene ID | Variant and effect |
| --- | --- | --- | --- | --- |
| INV271 E6 | 14 | 724332 | EPAC (PF3D7_1417400) | Indel frameshift |
| INV293 G4 | 14 | 721293 | EPAC (PF3D7_1417400) | Indel frameshift |
| INV296 G1 | 14 | 725393 | EPAC (PF3D7_1417400) | Premature stop codon K3366* |
| INV280 C8 | 14 | 719556 | EPAC (PF3D7_1417400) | Premature stop codon E1422* |
| INV293 F10 | 13 | 853449 | Unknown function (PF3D7_1320700) | Non-Synonymous Q1263R |

### Identification of sibling parasites in clones of a clinical isolate

To determine the relatedness between clones generated from the same isolate, the level of genomic Identity By Descent (IBD) was calculated [17], which considers the segments of chromosomes shared between haploid genomes. As expected, most clones from the same isolate had virtually identical genome sequences, reflecting clones of the same common genotype within an isolate. More interestingly, Clone E5 from isolate line 289 showed 60.8% IBD with the other two clones that were derived at the same time (Fig 6B), and with the majority genome in the bulk culture of the isolate at day 78 and 153. Based on allele frequencies in the bulk isolate sequence (Supplementary Fig. S1), the proportion of Clone 3 was approximately 5% at day 78 of culture, prior to cloning. This clone is a sibling of the majority genome, with 25 recombination breakpoints observed between them, similar to the average number among progeny from an experimental cross [21]. This is within the wider range of relatedness among parasites within infections as estimated by single-cell sequencing [7] or multi-locus genomic analysis of cloned parasites from clinical infections elsewhere in Africa [22].

## DISCUSSION

These results which focus mostly on multiple-genotype isolates extend the previous findings from single-genotype isolates, showing that loss-of-function mutants appear over time in a proportion of isolates. Overall, eight (29 %) of the 28 isolates analysed had a premature stop codon or frameshift mutant arising and significantly increasing in frequency. All these isolates were cultured in the same laboratory, and there was no significant difference between single-genome and multiple-genome isolates in the proportions affected. The proportion affected is also not significantly different from that previously seen in an analysis of single-genome isolates cultured in a different laboratory, in The Gambia [3]. The variants were first detected after different lengths of time in culture, but only very rarely within the first two months. If future studies on clinical isolates are to avoid the potential of phenotypes being affected by mutants emerging in culture, it may be preferable to focus on analysing parasites that have not been in culture for longer than approximately this length of time. In many cases, isolates that have been cultured for longer may still be mutant-free, but genome sequencing to screen for loss-of-function mutants may be important as a quality control if longer-term cultures are analysed.

Most of the multiple-genotype isolates showed overall reduction of genetic diversity over culture time, which is not explained by drug resistance allele frequencies nor loss-of-function mutations. Aside from genomic differences between parasites, it is likely that growth rates are modified by changes at an epigenetic level, and these may occur at different timepoints in different lineages. Previous analysis of exponential parasite multiplication rates in some of the isolates analysed here showed that the rates usually increased over time in culture, and that this was not associated with whether isolates had single or multiple genotypes [4].

Loss-of-function mutations in *EPAC* have previously been shown in long-term laboratory-adapted parasite lines [3], and experimental analysis has confirmed that the gene is not functionally involved in cyclic AMP signalling as had been previously considered [23]. Although such mutants have been detected in clinical isolates, it is interesting that most isolates still do not carry such mutation even after up to 7 months in culture. The *P. falciparum* mutation rate is high enough that, in theory, every individual nucleotide is likely to be mutated in a continuous culture flask[1, 2]. Two non-mutually exclusive hypotheses can reconcile these facts. Firstly, any individual novel variant that confers a small survival advantage is likely to be randomly lost from a population shortly after arising [24], in which case a longer culture adaptation might lead to more lines having time to acquire a loss-of-function mutation in *EPAC*, as seen in many long-term laboratory-adapted strains3. Secondly, it is possible that such mutations in *EPAC* would only confer a fitness advantage in the appropriate genome background, in which case some lines would never acquire a loss-of-function mutation in *EPAC*. It should be noted that bulk sequencing of multiple-genotype isolates is not usually sufficient for identifying whether loss-of-function mutants are under selection, as it may not be possible to separate the effect of such a variant from other genomic differences.

Acquiring drug resistance may be at the expense of fitness in an environment where the drug is absent, but the relationship between resistance alleles and fitness is not straightforward, particularly as isolates that contain resistance alleles may also have inherited compensatory mutations elsewhere in the genome. Evidence that drug resistance has a fitness cost is illustrated by chloroquine resistance in Africa, where chloroquine use declined after resistance reached high frequency, following which the *crt* K76T resistance allele frequency declined in many populations, indicating a fitness cost in the absence of drug selection [25]. *In vitro*, fitness costs may be detected by differential growth rates, which is subject to experimental conditions and depends also on the genomic backgrounds of the parasites, as illustrated by independent studies on variants of the *mdr1* gene [26, 27]. It is becoming increasingly clear that epistasis between variants of different genes contribute to parasite fitness, particularly in the context of drug resistance selection and associated fitness costs [28, 29], but also due to selective processes that are less well known [29].

Within a culture of any multiple genotype infection isolate, all parasite genotypes are subject to the same conditions, making this an appropriate setting for studying the genetic basis of growth competition. As illustrated here, faster growing lines tend to outcompete others, but the interaction is not necessarily linear. Generating data with larger sample sizes may enable future genome-wide association studies, by comparing slow growing genomes versus faster growing genomes and identifying allele frequencies significantly associated with the phenotype. R, adjusting culture parameters such as temperature [5], static or shaking conditions [30], erythrocyte blood group types [31], or nutrient concentrations [32], may be performed in future to test whether there may be selection for parasite variants suited to particular conditions.

Further, as illustrated by the demonstration of genetically-related clones within one of the isolate lines analysed here, analysing related clones from within infections may be an approach to help associate genetic loci to phenotypes, in a manner similar to a Quantitative Trait Loci analysis from a genetic cross [33].

## Ethical Approval

Approval for the sampling of blood from patients for parasite culture was granted by the Ethics committees of the Ghana Health Service, the Noguchi Memorial Institute for Medical Research at the University of Ghana, the Navrongo Health Research Centre, the National Ethics Committee for Health Research in the Republic of Guinea, the Ministry of Health in Senegal, the Ministry of Health in Mali, and the London School of Hygiene and Tropical Medicine, UK. Written informed consent was obtained from parents or other legal guardians of all participating children with malaria from whom blood samples were obtained, and additional assent was received from the children themselves if they were 10 years of age or older.

## Sequence data

The newly described parasite genome sequence data have been deposited in the European Nucleotide Archive, and accession numbers are all listed in Supplementary Table S1.

## Author Contributions

Conceptualisation by AC, LBS, AN-G, GAA, DJC; Data Curation by LBS, DPK, DJC; Formal Analysis by AC, LBS, DJC; Investigation by AC, LBS, ED, ADA, AN-G, MD, DPK, GAA, DJC; Writing of original draft by AC, LBS, DJC; Reviewing and editing of manuscript by AC, LBS, AN-G, GAA, DJC.

## Acknowledgements

This study was supported by funding from the European Research Council (AdG-2011-294428), the Royal Society (AA110050) and the UK Medical Research Council (MR/S009760/1). Support for AC was provided by a joint MRC Gambia-LSHTM fellowship. Genome sequencing was conducted at the Wellcome Sanger Institute, coordinated by the MalariaGEN Resource Centre with administration by Sonia Goncalves, Kim Johnson, and Vikki Simpson, with funding from The Wellcome Trust (grants 098051, 206194, 090770). We are grateful to all patients who participated in the study, and to staff of the Navrongo Health Research Centre in Ghana, National Institute for Public Health in Guinea, at Nioro du Sahel Health Centre in Mali, and at Pikine Health Centre in Senegal for support with initial sample collection and storage. We thank Inayat Bhardwaj and Steve Schaffner for contributing to graphical designs of Fig. 1 and Fig. 6B, respectively.

## References

1. Claessens A, Hamilton WL, Kekre M, Otto TD, Faizullabhoy A et al. Generation of antigenic diversity in Plasmodium falciparum by structured rearrangement of Var genes during mitosis. PLoS Genet 2014;10:e1004812.

2. Hamilton WL, Claessens A, Otto TD, Kekre M, Fairhurst RM et al. Extreme mutation bias and high AT content in Plasmodium falciparum. Nucleic Acids Res 2017;45:1889–1901.

3. Claessens A, Affara M, Assefa SA, Kwiatkowski DP, Conway DJ. Culture adaptation of malaria parasites selects for convergent loss-of-function mutants. Sci Rep 2017;7:41303.

4. Stewart LB, Diaz-Ingelmo O, Claessens A, Abugri J, Pearson RD et al. Intrinsic multiplication rate variation and plasticity of human blood stage malaria parasites. Commun Biol 2020;3:624.

5. Tinto-Font E, Michel-Todo L, Russell TJ, Casas-Vila N, Conway DJ et al. A heat-shock response regulated by the PfAP2-HS transcription factor protects human malaria parasites from febrile temperatures. Nat Microbiol 2021;6:1163-+.

6. MalariaGen, Ahouidi A, Ali M, Almagro-Garcia J, Amambua-Ngwa A et al. An open dataset of Plasmodium falciparum genome variation in 7,000 worldwide samples. Wellcome Open Res 2021;6:42.

7. Nkhoma SC, Trevino SG, Gorena KM, Nair S, Khoswe S et al. Co-transmission of related malaria parasite lineages shapes within-host parasite diversity. Cell Host Microbe 2020;27:93–103 e104.

8. Wong W, Griggs AD, Daniels RF, Schaffner SF, Ndiaye D et al. Genetic relatedness analysis reveals the cotransmission of genetically related Plasmodium falciparum parasites in Thies, Senegal. Genome Med 2017;9:5.

9. Duffy CW, Amambua-Ngwa A, Ahouidi AD, Diakite M, Awandare GA et al. Multi-population genomic analysis of malaria parasites indicates local selection and differentiation at the gdv1 locus regulating sexual development. Sci Rep 2018;8:15763.

10. Duffy CW, Assefa SA, Abugri J, Amoako N, Owusu-Agyei S et al. Comparison of genomic signatures of selection on Plasmodium falciparum between different regions of a country with high malaria endemicity. BMC Genomics 2015;16:527.

11. Trager W, Jensen JB. Human malaria parasites in continuous culture. Science 1976;193:673–675.

12. Oyola SO, Ariani CV, Hamilton WL, Kekre M, Amenga-Etego LN et al. Whole genome sequencing of Plasmodium falciparum from dried blood spots using selective whole genome amplification. Malar J 2016;15:597.

13. Bohme U, Otto TD, Sanders M, Newbold CI, Berriman M. Progression of the canonical reference malaria parasite genome from 2002-2019. Wellcome Open Res 2019;4:58.

14. Auburn S, Campino S, Miotto O, Djimde AA, Zongo I et al. Characterization of within-host Plasmodium falciparum diversity using next-generation sequence data. PLoS One 2012;7:e32891.

15. Manske M, Miotto O, Campino S, Auburn S, Almagro-Garcia J et al. Analysis of Plasmodium falciparum diversity in natural infections by deep sequencing. Nature 2012;487:375–379.

16. Fiume M, Williams V, Brook A, Brudno M. Savant: genome browser for high-throughput sequencing data. Bioinformatics 2010;26:1938–1944.

17. Schaffner SF, Taylor AR, Wong W, Wirth DF, Neafsey DE. hmmIBD: software to infer pairwise identity by descent between haploid genotypes. Malar J 2018;17:196.

18. Bushell E, Gomes AR, Sanderson T, Anar B, Girling G et al. Functional Profiling of a Plasmodium Genome Reveals an Abundance of Essential Genes. Cell 2017;170:260–272 e268.

19. Zhang M, Wang C, Otto TD, Oberstaller J, Liao X et al. Uncovering the essential genes of the human malaria parasite Plasmodium falciparum by saturation mutagenesis. Science 2018;360.

20. Sargeant TJ, Marti M, Caler E, Carlton JM, Simpson K et al. Lineage-specific expansion of proteins exported to erythrocytes in malaria parasites. Genome Biol 2006;7:R12.

21. Jiang H, Li N, Gopalan V, Zilversmit MM, Varma S et al. High recombination rates and hotspots in a Plasmodium falciparum genetic cross. Genome Biol 2011;12:R33.

22. Nkhoma SC, Nair S, Cheeseman IH, Rohr-Allegrini C, Singlam S et al. Close kinship within multiple-genotype malaria parasite infections. Proc Biol Sci 2012;279:2589–2598.

23. Patel A, Perrin AJ, Flynn HR, Bisson C, Withers-Martinez C et al. Cyclic AMP signalling controls key components of malaria parasite host cell invasion machinery. PLoS Biol 2019;17:e3000264.

24. Hastings IM. The origins of antimalarial drug resistance. Trends Parasitol 2004;20:512–518.

25. Rosenthal PJ. The interplay between drug resistance and fitness in malaria parasites. Mol Microbiol 2013;89:1025–1038.

26. Froberg G, Ferreira PE, Martensson A, Ali A, Bjorkman A et al. Assessing the cost-benefit effect of a Plasmodium falciparum drug resistance mutation on parasite growth in vitro. Antimicrob Agents Chemother 2013;57:887–892.

27. Hayward R, Saliba KJ, Kirk K. pfmdr1 mutations associated with chloroquine resistance incur a fitness cost in Plasmodium falciparum. Mol Microbiol 2005;55:1285–1295.

28. Amambua-Ngwa A, Button-Simons KA, Li X, al. e. The amino acid transporter pfaat1 modulates chloroquine resistance and fitness in malaria parasites. BioRxiv doi:101101/20220526493611 2022.

29. Band G, Leffler EM, Jallow M, Sisay-Joof F, Ndila CM et al. Malaria protection due to sickle haemoglobin depends on parasite genotype. Nature 2021.

30. Nyarko PB, Tarr SJ, Aniweh Y, Stewart LB, Conway DJ et al. Investigating a Plasmodium falciparum erythrocyte invasion phenotype switch at the whole transcriptome level. Sci Rep 2020;10:245.

31. Theron M, Cross N, Cawkill P, Bustamante LY, Rayner JC. An in vitro erythrocyte preference assay reveals that Plasmodium falciparum parasites prefer Type O over Type A erythrocytes. Sci Rep 2018;8:8133.

32. Kumar S, Li X, McDew-White M, Reyes A, Delgado E et al. A malaria parasite cross reveals genetic determinants of Plasmodium falciparum growth in different culture media. Front Cell Infect Microbiol 2022;12:878496.

33. Reilly Ayala HB, Wacker MA, Siwo G, Ferdig MT. Quantitative trait loci mapping reveals candidate pathways regulating cell cycle duration in Plasmodium falciparum. BMC Genomics 2010;11:577.

