## Supplementary for "Genomic variation during culture-adaptation of genetically complex *Plasmodium falciparum* clinical isolates"

**Supplementary Table S1.** Sources, accession numbers, read depth and $F_{WS}$ values of genomes of all isolates at each sequenced timepoint (Separate EXCEL file).

**Supplementary Table S2.** *De novo* mutations identified in *P. falciparum* clones derived from clinical isolates.

| Clone | Chr | Pos | Ref | Alt | Type | AA_CHANGE | EFFECT | GENE |
|---|---|---|---|---|---|---|---|---|
| **Line 280 C8** | 08 | 990744 | T | G | snp | D93E | NonSyn | PF3D7_0822400, Unknown function |
| **Line 280 F10** | 12 | 199488 | T | C | snp | R918G | NonSyn | PF3D7_1204100, Unknown function |
| **Line 286 B10** | 14 | 1582671 | A | C | snp | K641T | NonSyn | PF3D7_1439100, DEAD/DEAH box helicase |
| **Line 286 F1** | 04 | 672774 | T | G | snp | K2122T | NonSyn | PF3D7_0415200, Unknown function |
| **Line 286 F1** | 06 | 644353 | G | A | snp | S101N | NonSyn | PF3D7_0615500, CRK5 |
| **Line 278 B10** | 02 | 728797 | GAATA TATA | G | indel | . | INTRON | PF3D7_0217600, Unknown function |

**Supplementary Fig. S1.** Genome-wide plots of single nucleotide polymorphism allele frequencies within *P. falciparum* clinical isolates during culture adaptation. All isolates are shown, with plots as described in the legend of Fig. 3.

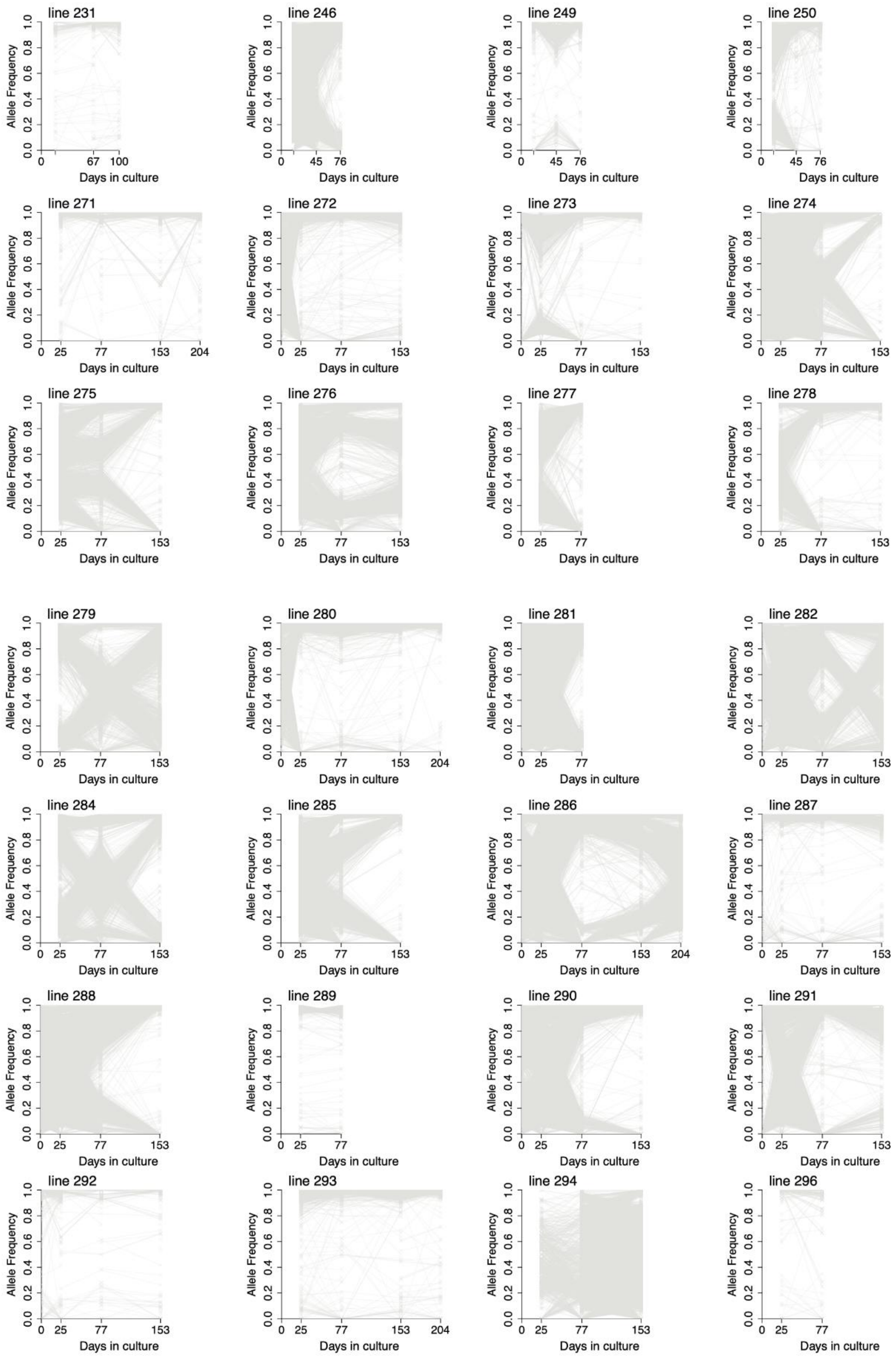

**Supplementary Fig. 2.** Allele frequencies at SNPs in *P. falciparum* drug resistance genes *dhfr*, *dhps* and *mdr1*, additional to those shown in Fig. 4, in multiple-clone isolates after different lengths of time in culture. For each polymorphism, isolates shown are only those that had mixed alleles at one or more timepoints. Across all isolates, there was no significant directionality to the changes in frequencies of any of these resistance-associated polymorphisms over time in culture.

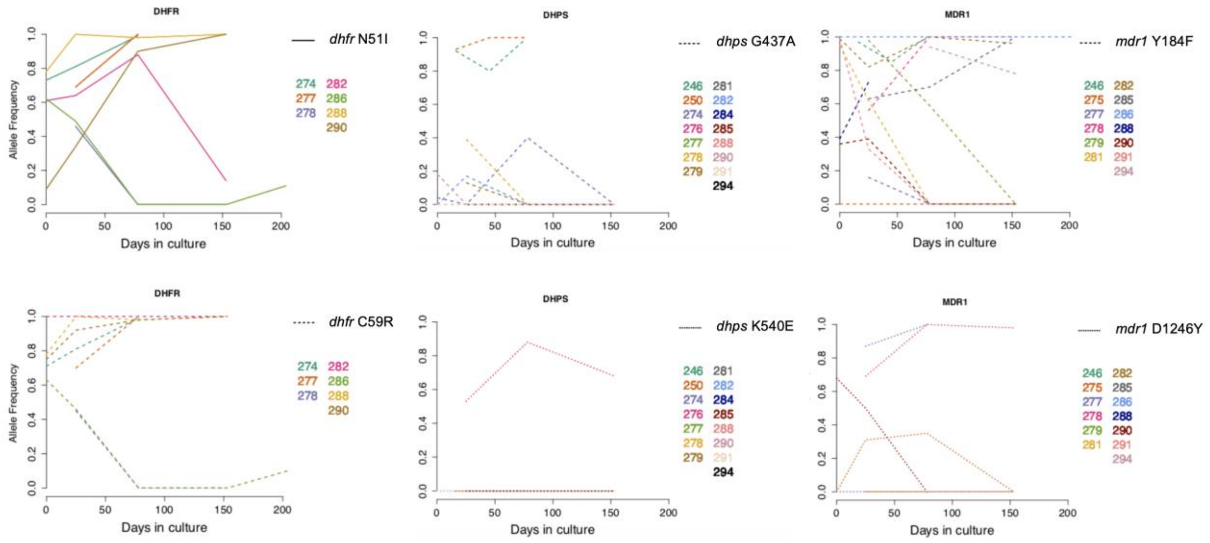
